# Temporal shift in task factor influence across the stretch reflex

**DOI:** 10.1101/2024.12.07.627335

**Authors:** Frida Torell, Robin Rohlén, Michael Dimitriou

## Abstract

Mechanical perturbations applied to the arm can elicit reflexive actions. These rapid corrective responses include the stretch reflex, which consists of different components: the short-latency reflex (SLR) as well as the early and late long-latency reflex (LLR). In this study, we explore how different task factors dynamically influence these reflex components in the context of a specific delayed-reach paradigm. Using multiple linear regression (MLR), we analysed reflex responses from seven muscles actuating the right arm to examine the effects of mechanical load, preparatory delay, perturbation and target direction, as well as their two-factor interactions. The MLR analysis shows that our delayed-reach tasks engaged shoulder girdle muscles, whereas the biceps and triceps primarily acted as stabilizing muscles, with rapid responses triggered regardless of perturbation direction. Specifically, our analyses show that the earliest corrective response, the SLR, exhibited some task/target-dependent modulation particularly in muscles of the shoulder girdle, although background (pre-)loading decreased this modulation. The SLR was primarily influenced by the main factors *Load* and *Perturbation*, along with the interaction *Load × Perturbation*. Perturbations aligned with the load direction were associated with increased electromyographic (EMG) activity across all examined muscles. While there was a small but significant effect of load during the early LLR, this effect diminished by the late LLR epoch. Task-dependent modulation was most pronounced at the late LLR epoch, suggesting greater top-down modulation of this reflex component. In particular, the late LLR was shaped by the factors *Perturbation* and *Target*, as well as the interaction *Perturbation × Target*. Targets and perturbations in opposing directions resulted in heightened EMG activity, and shoulder muscles exhibited stronger LLR responses for targets located farther along the muscle shortening direction. Across all reflex epochs, a longer preparatory delay (750 ms) was associated with higher EMG activity.

## Introduction

Accurate control of the arm is facilitated by corrective reflex actions. A prominent source of corrective actions is the so-called muscle stretch reflex. Such reflexive muscle contractions occur on a fixed time scale. They are divided into the Short Latency Reflex response (SLR) beginning approximately 20-25 ms after stretch onset, and the Long Latency Reflex (LLR) beginning 50 ms after stretch onset (Hammond, 1956; Marsden *et al*., 1972). At the level of the upper arm, muscle contractile activity appearing ∼100 ms after a mechanical perturbation is considered to be under voluntary control e.g., (Yang *et al*., 2011). The muscle stretch reflex involves short-loop (spinal) and long-loop neural pathways. Whereas in its simplest form, the SLR depends on spinal mechanisms, the LLR is thought to consist of both spinal and supraspinal components (Kurtzer, 2014; Soteropoulos & Baker, 2020). That is, the LLR responses (early and long, i.e. 50-75 and 75-100 ms post perturbation, respectively) are multifaceted and likely depend on multiple neural pathways. The neural circuits involved in LLRs likely allow interactions and integration of information from the spine, brainstem, and brain areas, including the cerebellum, posterior parietal cortex and frontal cortex (Pruszynski *et al*., 2011a; Pruszynski *et al*., 2011b; Pruszynski & Scott 2012).

It is well established that stretch reflex responses can allow for effective resistance against unwanted postural disturbances (Nichols & Houk, 1976). LLRs have been found to depend on verbal information and target location (Capaday *et al*., 1994; Crago *et al*., 1976; Lewis *et al*., 2006, Pruszynski *et al*., 2008; Rothwell *et al*., 1980). Moreover, in a bimanual postural task, unimanual reflex responses were modulated online within a single trial based on a complex interaction across the arms (Dimitriou *et al*., 2012). In addition to facilitating postural maintenance, reflex responses can also be modulated to accommodate the execution of movements in a task-dependent manner. For instance, in experiments involving active movement, the stretch reflex is significantly modulated in response to target shape (Nashed *et al*., 2012), obstacles (Nashed *et al*., 2014), static and moving targets (Cluff & Scott, 2015). For review, see (Scott, 2016). Moreover, there has been evidence that stretch reflex gains reflect the degree of visuomotor adaptation and the relative congruence between visual and haptic coordinate frames (Dimitriou, 2018).

Previous work examining reflex responses in the context of delayed reaching has demonstrated goal-directed tuning of the SLR and the LLR, with the latter being more robust against background muscle loading (Papaioannou & Dimitriou, 2021). In addition, we have shown that assistive loading (i.e., imposed unloading) facilitates goal-directed tuning of stretch reflexes at the shoulder (anterior and posterior deltoid as well as pectoralis) and that goal-directed tuning is less pronounced in the seemingly less dexterous non-dominant arm (Torell *et al*., 2023a; Torell *et al*., 2023b). Concerning the LLR, we have shown that there is an impact of different task factors (i.e., background load, perturbation direction, preparatory delay length and target direction). The aim of the present paper was to study in more detail the impact of these task factors on stretch reflex responses from the dominant arm, and determine the degree to which these factors interact and possibly vary across muscles actuating the upper limb (i.e., brachioradialis, biceps, lateral triceps, long triceps, anterior deltoid, posterior deltoid, and pectoralis). Specifically, we used a Multiple Linear Regression (MLR) approach to investigate how the different components involved in a delayed-reach task together impacted SLR and LLR responses across major muscles actuating the upper limb. Examining how factor influence shifts across reflex epochs and how our experimental design affects this temporal progression will enhance our understanding of the stretch reflex and inform the development of future studies.

## Materials and methods

### Subjects

This paper uses datasets from previously published studies (Torell, 2023; Torell *et al*., 2023a). The current study included data from 16 participants (eight males and eight females, mean age 26.9 ± 5.4 years) who performed Experiment ‘1’ and 16 participants (eight males and eight females, mean age 26.9 ± 5.4 years) who performed Experiment ‘2’. The inclusion criteria were (1) 18-40 years old, (2) 165-185 cm, (3) right-handed, (4) neurologically healthy, and (5) had normal movement range. All participants were naive to the purpose of the tasks prior to experimentation. The participants were financially compensated for their contribution and gave informed, written consent prior to participating in the study, per the Declaration of Helsinki. The current study forms part of a research program approved by the Ethics Committee of Umeå University, Umeå, Sweden. The dataset from Experiment 1 is available at Mendeley Data, V1, doi: 10.17632/44n4m42rvw.1. The Experiment 2 dataset is available at Mendeley Data, V1, doi: 10.17632/hnfp5yrght.1 (Torell *et al*., 2023a; Torell *et al*., 2023b).

### Robotic platform

In both experiments ‘1’ and ‘2’ (Torell, 2023; Torell *et al*., 2023a) participants were seated upright in an adjustable chair in front of the KINARM robotic platform (KINARM end-point robot, BKIN Technologies, Canada). The chair was adjusted to fit the height of each participant. The participants used their dominant right hand to grasp the robotic manipulandum (Figure 1A). The right forearm was placed inside a custom-made airsled that used compressed air to allow frictionless arm movement in a 2D plane. To ensure mechanical connection between the arm, the foam-cushioned airsled and the KINARM^TM^ handle, the forearm and hand were secured using a leather fabric with Velcro attachments. This attachment also fixated the wrist in alignment with the forearm. The angle of the wrist was 0° pronation/supination throughout the experiment. At the start position, the elbow was flexed by ∼90°. Once the participant was seated conformably, the chair was mechanically secured to the floor by four adjustable legs that also prevented chair rotation. During the experiment the participants rested their forehead against a cushioned and height-adjustable plate. In doing so the posture was further stabilized and the degree of neck fatigue was reduced. The robotic platform generates kinematic data of handle position and records the forces exerted by the participants’ hand, using a six-axis force transducer (Mini40-R, ATI Industrial Automation, USA) embedded in the robotic handle. Kinematic and force data from the robotic platform were sampled at 1 kHz.

**Figure 1.**
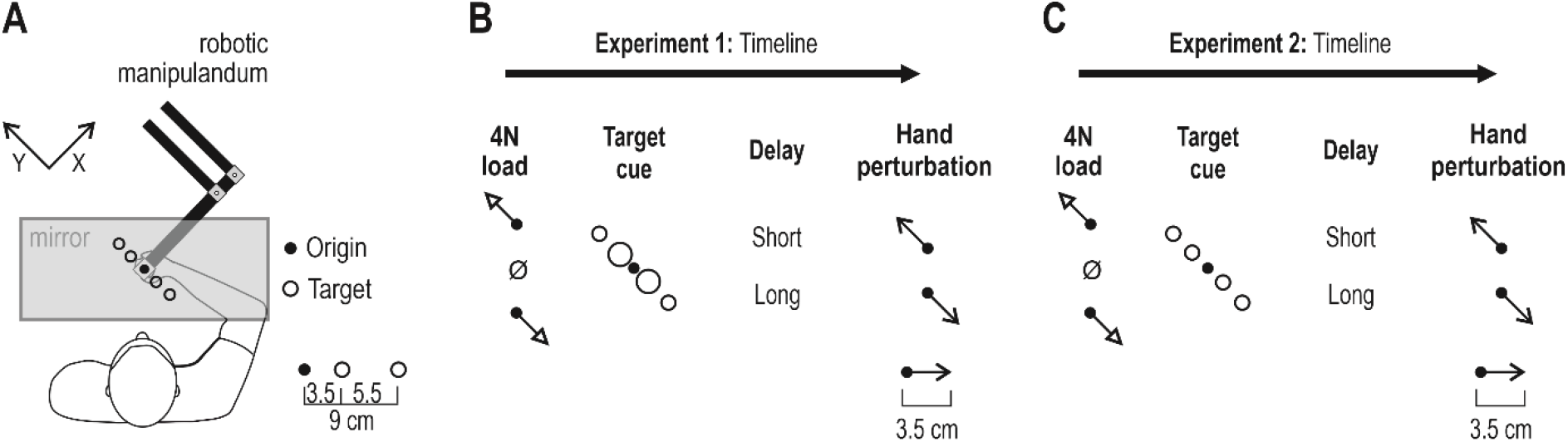
Experimental setup and behavioral tasks. **(A)** The participants were seated in front of the robotic manipulandum. Their dominant right hand was secured to the robot handle, and the forearm was resting on an airsled that allowed for frictionless movement on a 2D plane. The visual targets and origin were continuously displayed as orange circle outlines on a one-way mirror reflecting the projections of a monitor. The center of the handle was represented as a moving visual cursor. The participants started each trial by moving their hand to the origin location. **(B)** In Experiment ‘1’, the timeline consisted of four stages. After moving the cursor to the origin, one of three load conditions was applied (4N in -Y or + Y direction or null load). Then, a target was cued (turned red) for a long (750 ms) or short (250 ms) period, representing a preparatory delay. The same target then turned green (‘Go’ signal) and simultaneously the hand was perturbed by 3.5 cm in either the -Y or + Y direction, i.e., towards or in the opposite direction of the target. **(C)** Experiment ‘2’ differed from ‘1’ only in that all targets were of the same size.

### Experimental design

The experimental design of the delayed-reach tasks involving haptic perturbations is summarized in Figure 1 and described in more detail elsewhere (Torell, 2023; Torell *et al*., 2023a). Briefly, in both Experiments ‘1’ and ‘2’, a one-way mirror prevented a view of the hand and robotic handle, and on this mirror, visual stimuli were projected in the plane of movement. Visual stimuli included a moving white dot (‘cursor’; 1 cm diameter) tracking the position of the hand. In addition, four targets and the center point were continuously displayed as orange circle outlines (see, e.g., Figure 1B-C). The targets were placed in a front-and-left direction (defined as the ‘+Y’ direction) and right-and-back direction (‘-Y’ direction). The only difference between Experiments ‘1’ and ‘2’ was the size of the ‘near’ targets, i.e., all targets in Experiment ‘2’ had the same size (2.4 cm diameter). Each trial of either experiment started by moving the cursor to the origin circle.

For the trial to progress, the cursor had to remain immobile inside the origin for a random wait period ranging between 1 and 1.5 seconds. A 4N load was then applied in either -Y or +Y direction, or no load was applied. This stage of the task lasted 2 seconds (i.e., any load had an 800 ms rise-time and 1200 ms hold-time). One of the four targets was then cued by suddenly turning from an orange outline to a filled red circle. After a short or long preparatory delay (250 ms or 750 ms), a position-controlled perturbation of the hand was applied in the +Y or -Y direction (3.5 cm with 150 ms rise-time and no hold). The cursor position was frozen during the perturbation. The haptic perturbations were designed to produce approximately bell-shaped velocity profiles. At the onset of the perturbation, the filled red circle also turned green (referred to as the ‘Go’ signal). At the end of the perturbation, the participants were instructed to complete movement to the target, if necessary. After reaching the target and remaining there for 300 ms, the participants received visual feedback on their performance. ‘Too fast’ was shown if the time from the ‘Go’ signal to the time the target was reached was <400 ms, while ‘Correct’ was displayed if they reach the target between 400-1400 ms, and ‘Too Slow’ was shown if the target was reached after >1400 ms. After this the subsequent trial was started by moving the cursor to the origin.

The experiments represented a full factorial design without center points. A full factorial design enables the study of main effects and two-factor interactions. Specifically, each experiment consisted of 48 unique trials (i.e., experimental conditions): four targets (‘close’ and ‘far’ targets in both the -Y and the +Y direction) × three load conditions (no load or a 4N load applied in either the -Y or +Y direction) × two preparatory delay times (250 ms or 750 ms) × two perturbation directions (applied in either the -Y or +Y direction). One ‘block’ represents one set of the 48 unique trials. Within each block, the trials were randomized. Each experiment involved 15 blocks (i.e., 720 trials in total). The 15 replicates were collected since EMG data obtained with surface electrodes are known to be noisy (Alkan & Günay, 2012). If the participant needed to rest, they could pull the cursor to the side after any trial. However, breaks were encouraged to occur between blocks.

### Electromyography (EMG)

In the delayed-reach tasks from which the data in this study originated, surface EMG was recorded from the belly of (1) *m. brachioradialis* (BR), (2) *m. biceps brachii* (BB), (3) *m. triceps brachii caput laterale* (TriLat), (4) *m. triceps brachii caput longum* (TriLo), (5) *m. deltoideus pars anterior* (DeltA), (6) *m. deltoideus pars posterior* (DeltP), and (7) *m. pectoralis major* (PecM). We used Bagnoli^TM^ DE-2.1 electrodes (Delsys Inc., USA) with contact dimensions 10.0 × 1.0 mm and 10 mm inter-electrode spacing. Prior to attaching the EMG electrodes, the skin was cleaned using alcohol swabs and the electrodes coated with conductive gel. The electrodes were placed on the peak of the belly of the studied muscles in the direction of the muscle fibers. All electrodes were attached with double-sided tape and secured using surgical tape. One ground electrode (Dermatrode® HE-R Reference Electrode type 00200-3400; American Imex, Irvine, CA, USA), with a diameter of 5.08 cm, was placed on the *processus spinosus* of C7, secured using surgical tape, and connected to the same EMG system (Bangoli-8 EGM System, Delsys Inc., USA). The EMG signals were band-pass filtered online through the EMG system (20 – 450 Hz) and sampled at 1 kHz.

## Data preprocessing

The EMG data was high pass filtered using a fifth-order, zero phase-lag Butterworth filter with a 30 Hz cutoff. In addition, a rectification step was included to obtain the shape of the EMG signal. Normalization allowed EMG data from different muscles and participants to be compared and combined. The raw data was normalized (z-transformed) using a procedure described in detail elsewhere (Dimitriou, 2014; Dimitriou, 2016; Dimitriou, 2018). In summary, the procedure involves concatenating the EMG data for each participant and muscle individually to calculate a grand mean and grand standard deviation. Normalized EMG data was obtained by subtracting the grand mean of a specific muscle from the measured value and dividing it by the grand standard deviation of that muscle. The first five blocks of trials (i.e., repetitions of unique trial conditions) were viewed as part of a familiarization period and removed from further analyses. The normalized single trial data for the latter ten blocks were then averaged (mean) for each participant, muscle and experimental condition. These normalized averages were compared and a global mean across participants was also generated for each experimental condition.

## Statistical Analyses

Statistical analyses were done using the z-normalized EMG data, which was first rectified and filtered as described above. Preprocessing and great means were done using MATLAB® (MathWorks, Natick, MA, USA). Multiple linear regression was performed using MODDE®, version 11 (Sartorius, Umeå, Sweden).

### Multiple linear regression (MLR)

Multiple linear regression is a predictive modeling method used to determine the relationship between several x-variables and one or more y-variables (Fisher, 1922; Pearson, 1903; Yule, 1897). Regression is performed between **X** and **Y**, so that the sum of squares of the error term is minimized. Important to know is that MLR requires the data not to have co-linearity or inference between variables in **X, X** not to be noisy, and that the data set to have more samples than variables (Esbensen, 2002). Owing to these limitations, MLR is most suited in cases where the x-variables are controlled by a design of experiments (DOE). MODDE® (Sartorius, Umeå, Sweden) uses single value decomposition (SVD) to solve the **Y** = **B**\***X**+**E**, where **B** is the regression coefficient matrix, and **E** is the residual matrix (Eriksson *et al*., 2008). The equation is solved to minimise the least squares between observed and predicted values. In this way, MODDE uses SVD to derive a pseudo-inverse of **X**, to identify the best approximate solution in terms of least squares.

When fitting a regression model, the most important diagnostic tool consists of the two model estimates, R^2^ and Q^2^. The R^2^ parameter is referred to as *goodness of fit* and is a measure of how well the regression model fits the raw data. The R^2^ parameter varies between 0 and 1, where 1 indicates that the model fits the raw data perfectly and 0 that there is no model at all (Esbensen, 2002). The main disadvantage of R^2^ is that it comes closer and closer to 1 by including more model terms. Hence, we should not only inspect the R^2^ parameter to validate our models. If we combine the interpretation of the R^2^ parameter with the Q^2^ parameter called the *goodness of prediction*, we have a better indication of the usefulness of the model. The Q^2^ parameter provides a measure of the predictive power of the model. The Q^2^ parameter varies between minus infinity and 1. As a rule of thumb, a Q^2^ >0.5 should be regarded as good, and Q^2^ >0.9 as excellent, and both R^2^ and Q^2^ should be high, and preferably not separated by more than 0.3 (Esbensen, 2002). When the R^2^ and Q^2^ differ more, this is a sign that we have an unacceptable model.

In the present paper, MLR was used to study z-scored EMG activity observed during three time periods: the SLR period, as well as the early and late LLR. Each of these epochs was 25 ms long. Specifically, the haptic perturbation was viewed as the starting point (i.e., zero time = perturbation onset), SLR 25-49 ms, the early LLR 50-74 ms and late LLR 75-99 ms post perturbation onset. The different epochs are exemplified by a representative participant in Figure 2. The MLR analyses were performed using the average EMG activity of all participants across the five different epochs. In other words, the MLR models were calculated based on the grand mean across all participants. Since EMG data is noisy, averaging was done by obtaining the mean value to ensure the identification of the proper responses. Regression coefficients were calculated for four main task factors (load, preparatory delay, perturbation and target direction) and 44 two-factor interactions. The regression coefficients can be interpreted as mean values and the 95% confidence intervals as the precision of these means. Their size of the 95% confidence intervals depends on three factors: (1) the quality of the experimental design, (2) the goodness (R^2^) of the regression model, and (3) the number of degrees of freedom (Eriksson *et al*., 2008). A confidence interval that crosses zero is not statistically significant, as there is no consensus regarding the direction of this regression coefficient. Here, significant factors were determined using the 95% confidence interval.

**Figure 2.**
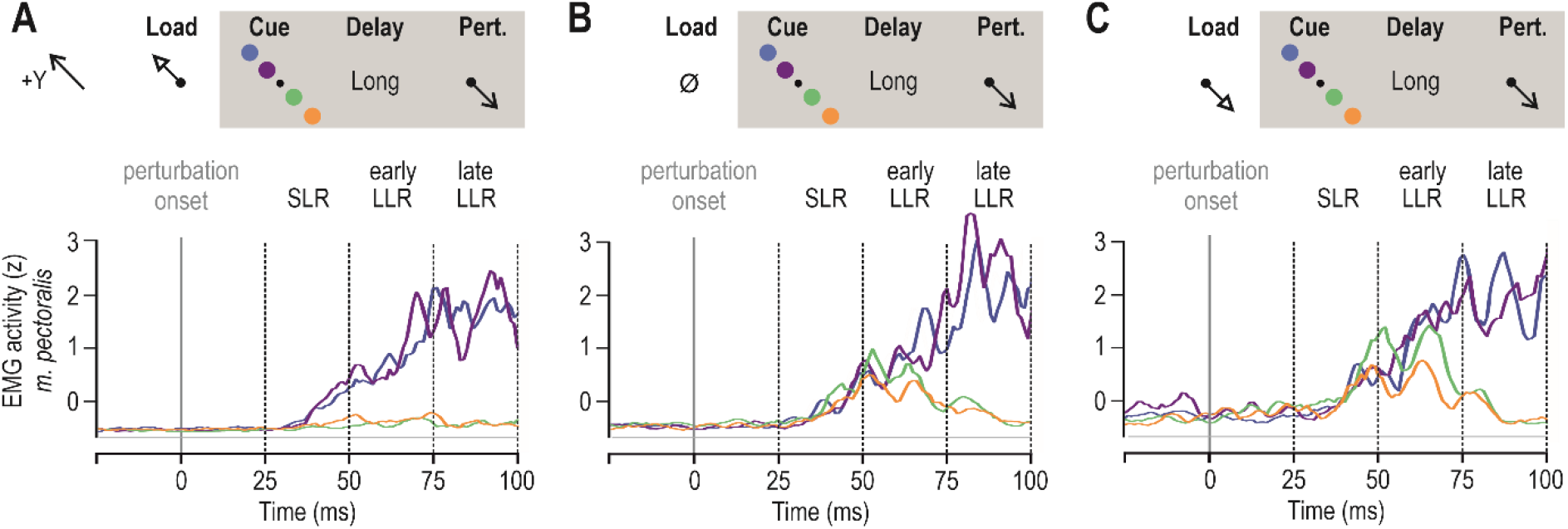
Averaged pectoralis EMG activity of an exemplary participant in Experiment ‘2’. The blue and purple traces represent ‘far’ and ‘near’ targets in the +Y direction, respectively. Reaching these targets required shortening of the pectoralis. The orange and green traces represent ‘far’ and ‘near’ targets in the -Y direction, respectively. These trials required a stretch of the pectoralis during reaching. All data in this figure involved a long preparatory delay (i.e., 750 ms) and are aligned on perturbation onset (time 0). **A**, A slow-rising load was first applied in the +Y direction (unloading the pectoralis) followed by a visual target cue and subsequently a haptic perturbation that stretched the pectoralis (see also Fig. 1A). **B**, As in A, but no slow-rising load was applied. **C**, As in A, but the slow-rising load was applied in the –Y direction, loading the pectoralis.

## Results

Figure 2 displays a representative example of pectoralis EMG activity across the different stretch reflex epochs induced in the delayed-reach task. The different panels (A-C) represent different background mechanical loads on the hand and coloring represents different visual targets. Reaching the blue and purple targets required shortening of the pectoralis while the green and orange targets required stretching of the pectoralis.

To explore the outcome of the experiments, each EMG trace was summarized according to the observed mean peak amplitude (one grand mean value for each examined epoch, i.e., SLR, early LLR and late LLR), and MLR models were created. The different trials represent the various combinations of factor settings that can be achieved and were used to study the five different time epochs. The design lacked center points but was run in 15 replicates per participant x 15 participants, so it was still possible to study the variability of the data. The MLR was done to study factor interactions between the four main effects (load, delay, perturbation, and target) and 44 two-factor interactions. The MLR models are summarized in Table 1.

**Table 1.** MLR model summary. Muscles included in the analysis: *m. brachioradialis* (BR), *m. biceps brachii* (BB), *m. triceps brachii caput laterale* (TriLat), *m. triceps brachii caput longum* (TriLo), *m. deltoideus pars anterior* (DeltA), *m. deltoideus pars posterior* (DeltP), and *m. pectoralis major* (PecM). Light grey numbers indicate that the difference between the R^2^ and the Q^2^ is greater than 0.3, i.e., that the model for this particular muscle was not statistically significant.

|  |  | SLR |  | Early LLR |  | Late LLR |  |
| --- | --- | --- | --- | --- | --- | --- | --- |
| | | $R^2$ | $Q^2$ | $R^2$ | $Q^2$ | $R^2$ | $Q^2$ |
| Experiment 1 | BR | 0.93 | 0.72 | 0.99 | 0.94 | 0.98 | 0.89 |
|  | BB | 0.74 | -0.12 | 0.93 | 0.71 | 0.95 | 0.78 |
|  | TriLat | 0.93 | 0.70 | 0.96 | 0.82 | 0.94 | 0.73 |
|  | TriLo | 0.94 | 0.75 | 0.96 | 0.84 | 0.98 | 0.93 |
|  | DeltA | 0.95 | 0.77 | 0.99 | 0.94 | 0.97 | 0.89 |
|  | DeltP | 0.99 | 0.97 | 0.99 | 0.96 | 0.99 | 0.97 |
|  | PecM | 0.98 | 0.90 | 0.99 | 0.96 | 0.98 | 0.93 |
| Experiment 2 | BR | 0.99 | 0.94 | 0.99 | 0.97 | 0.99 | 0.96 |
|  | BB | 0.82 | 0.22 | 0.93 | 0.68 | 0.97 | 0.85 |
|  | TriLat | 0.97 | 0.86 | 0.92 | 0.63 | 0.97 | 0.85 |
|  | TriLo | 0.98 | 0.91 | 0.97 | 0.89 | 0.99 | 0.95 |
|  | DeltA | 0.98 | 0.91 | 0.96 | 0.83 | 0.98 | 0.91 |
|  | DeltP | 0.99 | 0.96 | 0.99 | 0.97 | 0.99 | 0.98 |
|  | PecM | 0.99 | 0.94 | 0.98 | 0.90 | 0.99 | 0.95 |

As can be seen in Table 1, the MLR models across the stretch reflex epochs were best fitted and predicted for muscles of the shoulder girdle: anterior deltoid, posterior deltoid, and pectoralis. For these muscles, the range of the R^2^ values is extremely high, ranging from 0.95-0.99. Q^2^ values are also high, with a range of values 0.77-0.98. For the SLR epoch, the only weak model was for the biceps (BB; see Table 1). This was the case both for Experiment ‘1’ and ‘2’. The regression coefficients for the MLR model containing SLR data from Experiment 1 can be seen in Figure 3. The equivalent plot for Experiment 1 is available as Supplementary data (see Figure S1). The regression coefficients observed in Figures 3 and Supplementary Figure S1 show that EMG activity was impacted by the main factors ‘Load’, ‘Perturbation,’ and ‘Preparatory Delay’. Experiencing a perturbation in the -Y direction increased EMG activity in the biceps, lateral triceps, anterior deltoid and pectoralis. On the other hand, perturbations in the +Y direction positively affected the EMG activity of the brachioradialis, long triceps and posterior deltoid.

**Figure 3.**
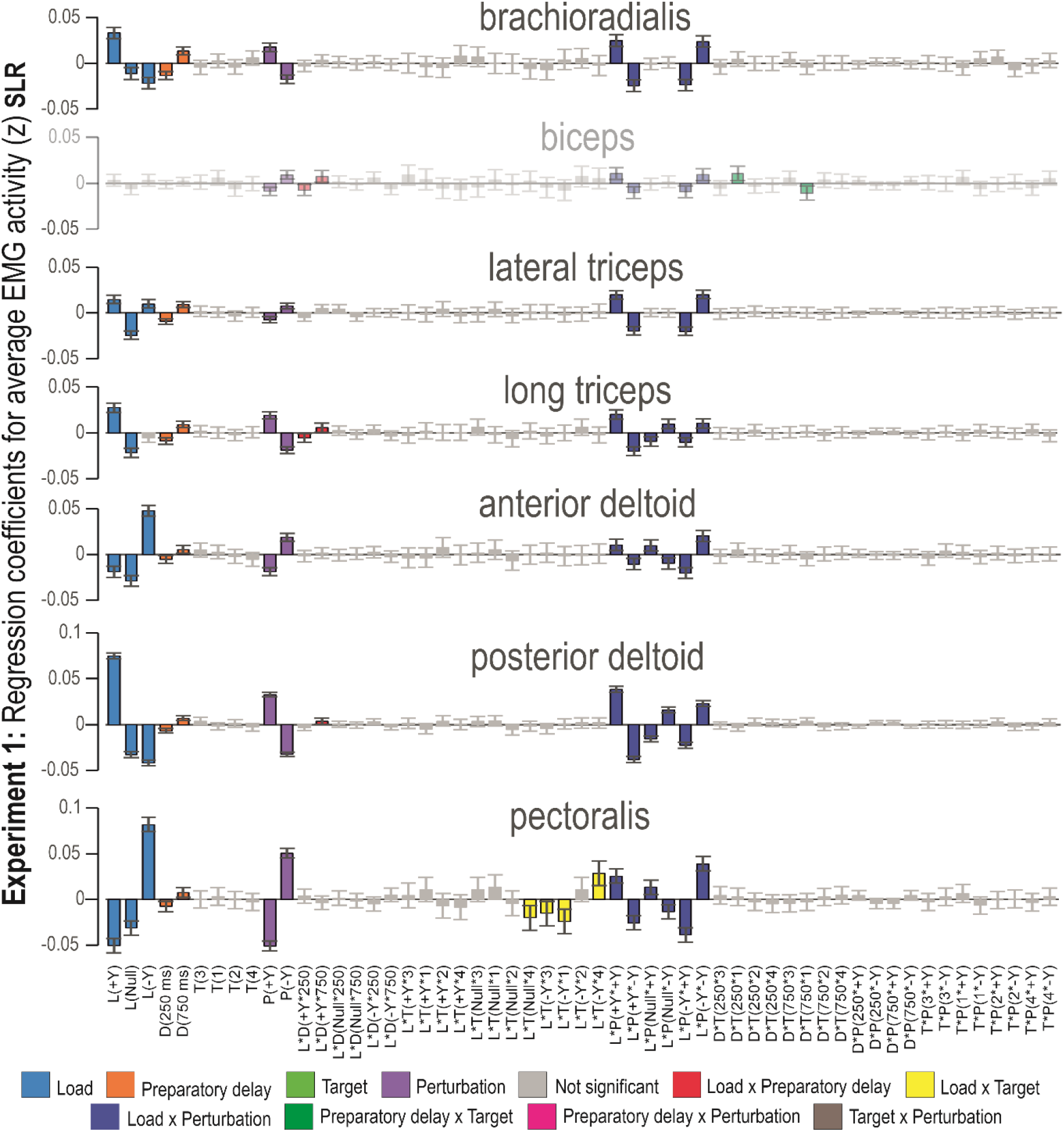
MLR coefficients for the SLR epoch of Experiment 1. The -Y and +Y directions are the same as defined in Figure 1. Coloring represents factor type; main factors and two-factor interactions whose 95% confidence interval includes zero are indicated as not significant (i.e., grey bars). The biceps bars are faded to indicate that the MLR model is not statistically significant. Abbreviations: L = load, D = preparatory delay, T = target, P = perturbation.

There was relatively increased EMG activity in trials where the perturbation was in the direction of muscle stretch. Short preparatory delays (250 ms) resulted in decreased EMG activity, while longer preparatory delays (750 ms) were associated with increased EMG activity. This was true for all studied muscles in both experiments, except for the biceps in Experiment 1. Specifically, the two-factor interaction ‘Load × Perturbation’ was significant for all studied muscles, where a perturbation in the load direction was associated with increased EMG activity for all included muscles. Regarding ‘Load × Perturbation’, the pectoralis and anterior deltoid displayed the opposite EMG pattern in response to the null load trials, compared to the pattern seen in the posterior deltoid and long triceps. For the pectoralis, the two-factor interaction ‘Load × Target’ showed that EMG activity was increased when the participant experienced a -Y load (i.e., after loading the pectoralis) and had a target in the -Y direction.

The regression coefficients for the MLR model containing early LLR data from Experiment 2 can be seen in Figure 4. The equivalent plot for Experiment 1 is available as Supplementary data. The regression coefficients illustrated in Figure 4 show that all main effects were significant for most of the investigated muscles. The perturbation effect was now the greatest, while the load effect was smaller. Regarding target direction, having targets in the +Y direction increased EMG activity in the pectoralis, anterior deltoid and lateral triceps, while targets in the -Y direction increased EMG activity of the posterior deltoid and long triceps. The early LLR was characterized by ‘Target’, where we see an overlap of the confidence intervals for targets in the same direction, meaning that the target direction was significant and that there was no effect of target distance in this epoch. The most important two-factor interactions during the early LLR epoch were ‘Load × Perturbation’ and ‘Target × Perturbation’. For all investigated muscles, experiencing a perturbation in the same direction as the load increased EMG activity. In contrast, trials where the participants experienced a perturbation in the opposite direction were associated with decreased EMG activity.

**Figure 4.**
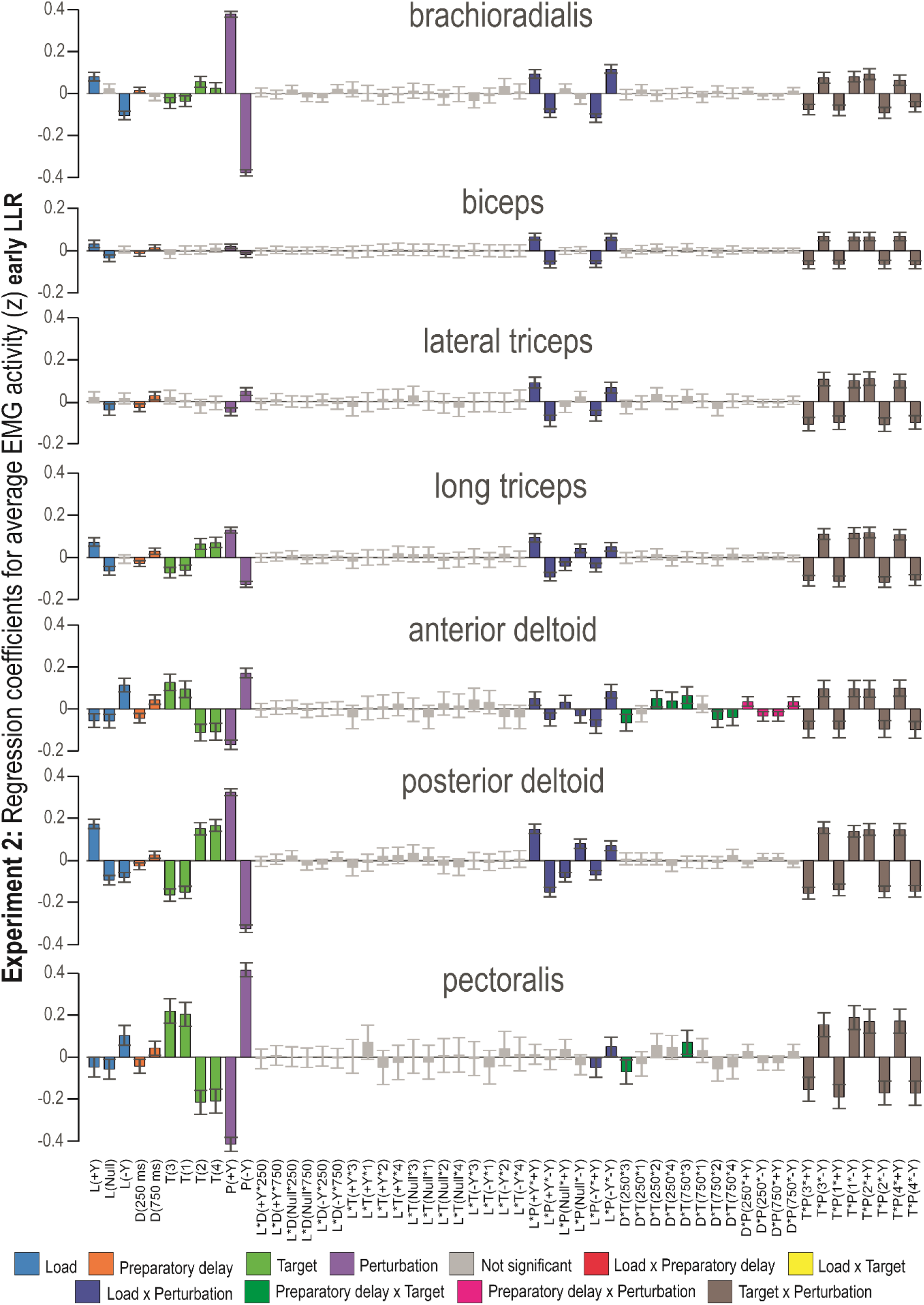
MLR coefficients for the early LLR epoch of Experiment 2. The -Y and +Y directions are the same as defined in Figure 1. Coloring represents factor type; main factors and two-factor interactions whose 95% confidence interval includes zero are indicated as not significant (i.e., grey bars). Abbreviations: L = load, D = preparatory delay, T = target, P = perturbation. The equivalent plot for Experiment 1 is available as Supplementary data.

In addition, the two-factor interaction ‘Target × Perturbation’ had a strong effect, with a similar pattern emerging for each of the studied muscles. Specifically, when experiencing a perturbation in the -Y direction, cued targets in the +Y direction were associated with increased EMG activity and vice versa. In other words, targets in the +Y direction and a perturbation in the +Y resulted in decreased EMG activity. In comparison, a perturbation in the -Y direction was associated with an increased EMG activity. The opposite was true for targets in the -Y direction, i.e., a perturbation in the +Y direction gave an increased EMG activity. This pattern was evident at the early LLR epoch across all studied muscles in both Experiment 1 and 2. For three muscles (i.e., anterior deltoid, posterior deltoid and brachioradialis), there was a small but significant effect of the two-factor interaction ‘Preparatory Delay × Perturbation’ in Experiment 1 and/or 2. For the two-factor interaction ‘Preparatory Delay × Perturbation’, a perturbation in the direction of muscle shortening was associated with an increased EMG activity. The shoulder muscles and long triceps also had a small but significant effect of ‘Preparatory delay × Target’.

The coefficients for factor interactions observed during the late LLR period of Experiment 2 are displayed in Figure 5. The equivalent plot for Experiment 1 is available as Supplementary data (Figure S3). The largest main factor effects are attributed to ‘Perturbation’ and ‘Target’. The effect of ‘Load’ is relatively smaller but still significant for some of the muscles investigated. ‘Preparatory Delay’ has a small but statistically significant impact on late LLR, with the longer preparatory delay inducing increased EMG activity. The only exception to this pattern is the brachioradialis, where the shorter delay was associated with increased EMG activity. For all investigated muscles, except for the biceps, ‘Target’ had a significant effect. The brachioradialis, long triceps and posterior deltoid displayed increased EMG activity when a target was in the -Y direction. For the pectoralis and anterior deltoid, increased EMG activity was seen when the target was in the +Y direction. In the direction of muscle shortening, the shoulder muscles (pectoralis, anterior deltoid, posterior deltoid and the long triceps) appear to have a significant difference attributed to target distance, indicated by the fact that the confidence intervals do not overlap. The increased EMG activity was recorded for the further targets in this epoch. The impact of the ‘Perturbation’ factor follows the same pattern as in the SLR epoch.

**Figure 5.**
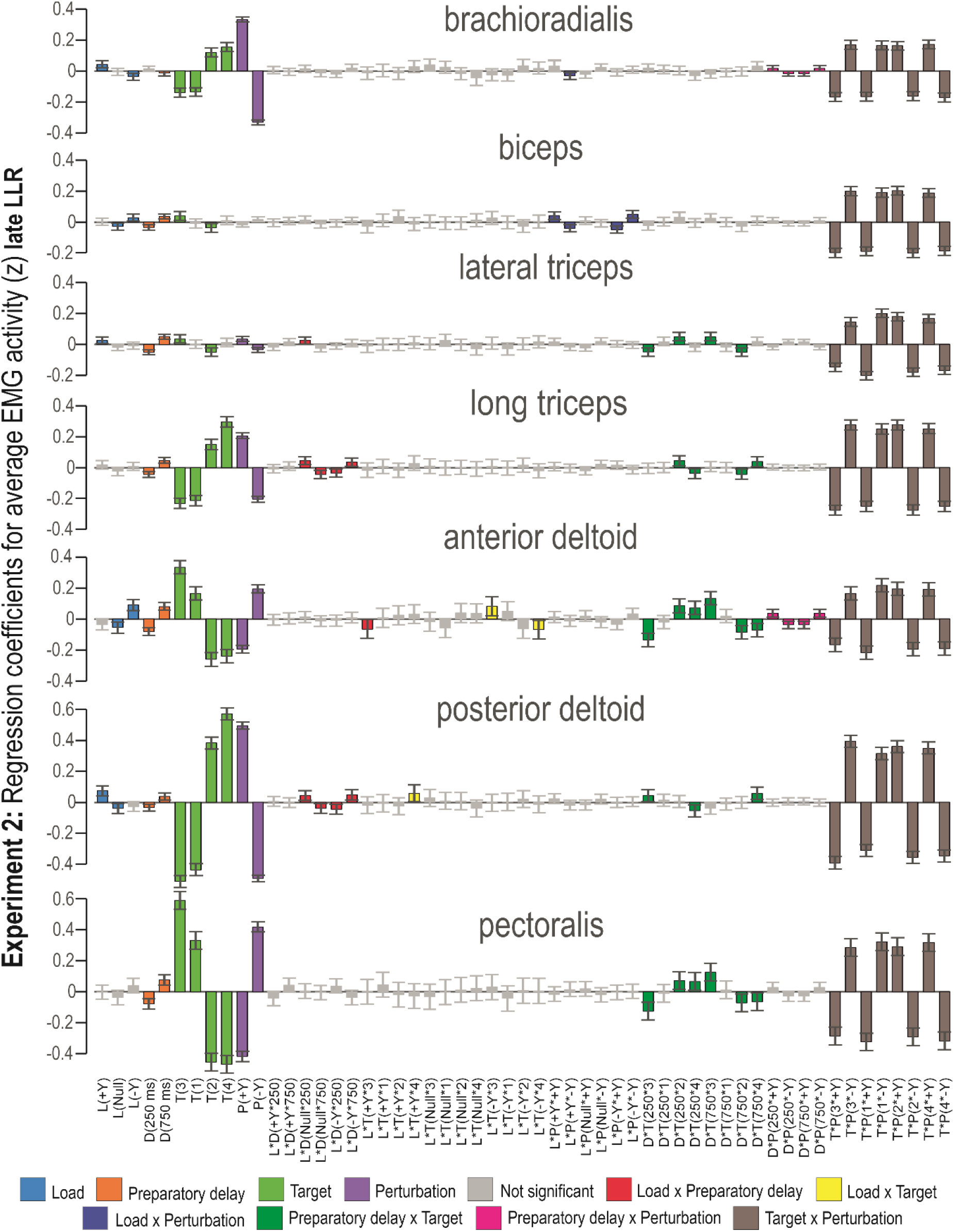
MLR coefficients for the late LLR epoch of Experiment 2. The -Y and +Y directions are the same as defined in Figure 1. Coloring represents factor type; main factors and two-factor interactions whose 95% confidence interval includes zero are indicated as not significant (i.e., grey bars). Abbreviations: L = load, D = preparatory delay, T = target, P = perturbation.

Compared to the early LLR epoch, the impact of ‘Load × Perturbation’ was smaller on late LLR but still significant for five muscles. In contrast, the two-factor interaction ‘Preparatory Delay × Target’ was significant in both the shoulder muscles and the two triceps muscles. In addition, the two-factor interaction ‘Preparatory Delay × Perturbation’ had a small but significant effect on EMG activity of the brachioradialis, long triceps and lateral triceps. Similar to what was seen during early LLR, the global pattern (observed in each EMG) shows that the two-factor interaction ‘Target × Perturbation’ was the same for all studied muscles, but the level of significance is higher in the shoulder muscles. This pattern indicates that targets and perturbations in the same direction are negatively correlated and that targets and perturbations in opposite directions resulted in increased reflex EMG. This pattern was seen for all studied muscles, despite the significance of the main factors ‘Target’ and ‘Perturbation’ (see, e.g., biceps and lateral triceps).

## Discussion

When the arm is perturbed, control is facilitated by corrective actions operating at different time scales. To understand how the influence of different factors changes over the stretch reflex timescales, an MLR analysis of EMG activity was employed in this study. With this approach, it was possible to probe how main factors and two-factor interactions concerning reach preparation time, background load, haptic perturbation and target direction contribute to limb control via their impact on stretch reflex responses. Using MLR allowed us to assess the combined reflex activity and interaction of multiple muscles as it unfolded across time and workspace. We found a clear shift in which factors contribute the most to the EMG activity across the stretch reflex epochs. The SLR displayed some task-dependent modulation, especially at the level of the shoulder muscles. There was a stronger task-dependent modulation of the late LLR response compared to the early LLR and SLR. In addition, we found a small but often significant effect of ‘Preparatory delay’, where a longer preparatory delay was associated with increased EMG activity. This was observed for all studied reflex epochs in both Experiment 1 and 2. In our delayed-reach tasks, we also found that the biceps and lateral triceps mainly functioned as stabilizing muscles, regardless of perturbation direction.

The SLR epoch was more strongly characterized by ‘Perturbation’ and ‘Load’, as well as the two-factor interaction ‘Perturbation × Load’. For shoulder muscles (i.e., anterior/posterior deltoid and pectoralis), we found that EMG activity during the SLR epoch is tightly coupled with perturbation direction and strongly influenced by the conditions experienced prior to perturbation (i.e., background load and preparation duration). Regarding the impact of load, this aligns with previous work by Pruszynski *et al*. showing that the SLR of upper extremities is characterized by ‘automatic’ gain-scaling (Pruszynski *et al*., 2011b). Moreover, our MLR analyses confirmed previous results showing that preparation duration impacts EMG activity at the SLR epoch, with longer preparatory delays (>250 ms) allowing appropriate goal-directed tuning of the response. In the context of postural maintenance, the reflex arc of the SLR is generally thought to represent a negative feedback response to local muscle stretch (Kurtzer, 2014), with the gain of this response being proportional to background loading (Matthews, 1986; Pruszynski *et al*., 2009). For Experiment ‘2’ in particular, there was also a small but significant impact of planned target direction on SLR magnitude, especially for the shoulder muscles. This is also in line with previous findings from our lab concerning stretch reflex responses (Papaioannou & Dimitriou, 2021; Torell *et al*., 2023a; Torell *et al*., 2023b). Such goal-directed tuning of SLR responses in delayed-reach suggests the involvement of independent fusimotor control of muscle spindles, as elaborated previously (Dimitriou, 2021; Dimitriou, 2022; Papaioannou & Dimitriou, 2021).

While the SLR of upper extremities is characterized by automatic gain-scaling, the LLR is robustly task-dependent (Pruszynski *et al*., 2011b). The early LLR has previously been shown to contain a stabilizing component modulated independently of voluntary action planned in relation to a cued target (Lee & Perreault, 2019). Both the early and the late LLR are said to have transcortical components where afferent feedback travels up the spinal cord to the motor cortex, eliciting motor commands that impact LLR responses (Phillips, 1969; Phillips *et al*., 1971). The late LLR occurs earlier than voluntary responses but is proposed to share some supraspinal pathways and functional capabilities associated with voluntary control (Kurtzer, 2014; Pruszynski *et al*., 2011b). This proposal also aligns with our current results, where we see more top-down modulation in the late LLR epoch. For example, we show that perturbation direction is a stronger contributor to EMG activity during the early LLR epoch, while target direction is a stronger contributor to EMG activity in the late LLR epoch (see Figures 4 and 5). Indeed, the late LLR has been previously found to display a high degree of task dependency (Lee & Perreault, 2019). This indicates that at least some components of the LLR response are highly adaptable and not strictly “reflex” in nature (Shemmell *et al*., 2010). Congruent with this notion is the belief that the primary motor cortex and reticular formation contribute to the LLR by generating or scaling to account for arm biomechanics, while the cerebellum scales the feedback response (Kurtzer 2014). It is evident that the late LLR contributes most in terms of facilitating the planned motor action via potent online corrections to haptic perturbations.

The results of the current study also support previous claims that multiple convergent pathways can contribute to the LLR (Kurtzer, 2014; Lee & Perreault, 2019; Lewis *et al*., 2004; Shemmell *et al*., 2010). There is evidence that, at least in part, the late LLR is mediated by motor cortical pathways (Lee & Hu, 2019). This is further strengthened by experiments where signals from the sensorimotor cortex have been blocked, resulting in diminished reflex gain modulation (Kimura *et al*., 2006). On the other hand, experiments with spinalized cats and monkeys indicate preservation of muscle activity in the LLR epoch (Ghez & Shinoda, 1978; Miller & Brooks, 1981). This suggests that transcortical pathways are not the only ones contributing to the LLR response, which likely results from a combination of multiple pathways, including transcortical and spinal neural circuits (Soteropoulos & Baker, 2020).

The specific delayed-reach paradigm investigated here primarily induced task-relevant stretch reflex responses in shoulder muscles (i.e., pectoralis and anterior and posterior deltoid). Shoulder muscles with a similar main axis of action (i.e., pectoralis and anterior deltoid) displayed a very similar reflex EMG pattern. However, a statistically significant MLR model was also generated for biceps LLR activity, involving a significant contribution from the two-factor interaction ‘Target × Perturbation’. As indicated in a previous study (Torell, 2023), out of the seven investigated muscles, the biceps contributed the least to muscle synergy models. This was attributed to the fact that the biceps muscle is not only responsible for elbow flexion but also involved in the outwards rotation of the forearm and contributes to shoulder stabilization (Landin *et al*., 2017). Here, we show that the main factors ‘Target’ and ‘Perturbation’ exert a very small or statistically insignificant effect on the early and late LLR response of both the biceps and lateral triceps, congruent with previous findings (Papaioannou & Dimitriou, 2021; Torell, 2023; Torell *et al*., 2023a; Torell *et al*., 2023b). Taken together, current and previous findings indicate that the biceps and lateral triceps were primarily engaged in limb stabilization, regardless of perturbation direction. Therefore, in future experiments using a similar design (e.g., the same load and perturbation directions) we do not expect a task-relevant modulation of biceps and triceps reflex activity. But our findings do confirm that the specific delayed-reach task employed in our studies promotes modulation of shoulder girdle stretch reflexes for task-relevant facilitation of reach.

Moreover, this study reinforces that MLR modeling can be a powerful tool for deciphering the concurrent role of multiple muscles engaged in a task where multiple factors can exert an impact and interact.

### Concluding remarks

To summarize, for the particular load and perturbation directions of the employed delayed-reach task, we found that:

- Background loading engages shoulder muscles rather than the biceps or triceps
- The biceps and triceps work as supportive or stabilizing muscles with rapid responses triggered regardless of perturbation direction
- There is task-dependent tuning of the SLR from shoulder muscles but such tuning decreases with background loading
- Although there is a small but significant effect of load at the early LLR, this effect diminishes during the late LLR epoch
- There is a stronger task-dependent modulation of the late LLR compared to the early LLR

## Additional Information

## Acknowledgments

The authors would like to thank Carola Hjältén for assistance with data collection, and Anders Bäckström for technical support. This work was supported by grants awarded to Dr. Michael Dimitriou by Hjärnfonden (FO2024-0425-HK-88) and the Swedish Research Council (project 2020-02140). The funders had no role in study design, data collection and analysis, decision to publish, or preparation of the manuscript.

## Ethical statement

All participants gave informed written consent before participating in the study and were financially compensated for their contributions.

## Author contributions

M.D. and F.T. conceptualized and designed the study, F.T. analyzed the data, M.D., R.R., and F.T. interpreted the results and wrote the manuscript.

